# Field-Based Characterization of Temporal Interference Stimulation: Beyond the Target-Centric Perspective

**DOI:** 10.64898/2026.09.16.752209

**Authors:** Tiandi Chen, Congcong Huo, Guangjan Shao, Zhanhong Cao, Chao Li, Jizhong Liu, Zengyong Li

## Abstract

**Purpose:** Temporal interference (TI) stimulation is a promising non-invasive technique for deep brain neuromodulation, yet accurately characterizing its complex and spatially distributed electric fields remains a fundamental challenge. This study proposes a computational framework that integrates topological analysis with mesh-aware metrics to overcome the limitations of conventional global field descriptors.

**Methods:** The method introduces three interconnected components: connected component analysis, which extracts spatially continuous field regions above thresholds; discrete ellipsoid modeling, which quantifies the geometric centroid, directional spread, and anisotropy of each region; and Local Moran’s I to identify robust spatial clusters. The framework was applied to TI envelope field simulations using a tetrahedral head model.

**Results:** The few largest connected components above an elevated intensity threshold collectively delineate the effective stimulation targets more reliably than isolated field peaks. The approach provides highly interpretable visual outputs that directly map field topology to brain anatomy, including layered component maps, discrete ellipsoids, and clustering maps of field extrema.

**Conclusion:** By offering a topologically coherent and visually intuitive analytical toolbox, this work advances the precision and interpretability of TI field assessment, supporting target localization, focality quantification, and parameter exploration in translational neuromodulation research.

## 1 Introduction

Transcranial electrical stimulation (TES) can transform neuroscience[1] and the treatment and rehabilitation of neurological disorders[2] due to its non-invasive and portable nature. TES involves the application of weak currents (usually 2 mA) to the scalp, thereby facilitating or inhibiting neural activity. Model-based simulations have demonstrated that conventional two-electrode TES is limited in its ability to achieve focused stimulation in deep brain regions, a consequence of the rapid spatial dispersion of electric fields and the low impedance of the scalp. Subsequently, high-definition electrode configurations were introduced to enhance focality and increase stimulation intensity; however, the effectiveness in targeting deep brain structures remains under debate[3].

The temporal interference (TI) stimulation, proposed by Grossman et al. in 2017[4], was based on the principle of linear superposition of electric fields. The paradigm involves the application of two high-frequency sinusoidal waveforms with slightly different frequencies, resulting in a low-frequency envelope capable of selective stimulation at deep brain targets. Subsequent animal experimentation provided evidence of the efficacy of TI stimulation in motor enhancement through stimulation of the motor cortex[5], as well as in the induction and suppression of epileptic activity via hippocampal stimulation[6, 7]. Electric field measurements conducted in human cadavers[8] and non-human primates[9] demonstrated the focal characteristics of TI fields, indirectly supporting the validity and predictive utility of computational modeling approaches. Recent human studies have indicated the potential of TI stimulation in cognitive enhancement through modulation of the hippocampus in healthy subjects[10], and in the improvement of motor function scores in patients with Parkinson’s disease[11]. TI stimulation can thus achieve precise neuromodulation of deep brain structures when optimal field focality is maintained.

To achieve improved focality, several parameter optimization strategies for temporal interference (TI) stimulation have been proposed[12–14]. Similar to approaches in transcranial alternating current stimulation (tACS)[15, 16], these methods are generally leadfield-based, optimizing combinations of stimulation parameters by defining metrics such as electric field intensity or focality within a target region, subject to a set of constraints, to identify optimal configurations. This approach, which determines stimulation targets by optimizing the leadfield while overlooking the overall distribution of the electric field, is termed the Target-Centric Perspective. Research on the characteristics of the resulting electric fields has largely remained secondary to optimization-focused research efforts typically addressing global field properties and target-region-specific features. For instance, focality is in some cases quantified as the cube root of the volume in which the field intensity exceeds 50% of the maximum within the target region[15]. As a result, seemingly simple questions—especially those posed by researchers without a background in optimization theory—often remain difficult to answer. These include questions such as: “Where is the target if two electrode pairs are arbitrarily placed?” or “What does the resulting field distribution look like?” Ultimately, such inquiries point toward a fundamental issue: the need for appropriate metrics to characterize the key features of electric field distributions.

Although current optimization strategies do not explicitly depend on the characteristics of electric field distributions, such dependence becomes necessary in more complex scenarios, where optimization methods requiring explicit analytical expressions become ineffective. This reliance is especially pronounced in multi-channel temporal interference (m-TI) stimulation, where electrode configuration optimization depends predominantly on intuitive insights into electric field distributions[17]. Another emerging trend involves the adoption of nonstandard montage configurations[18–20] and electrode arrays[21, 22], where field characteristics help reduce the search space during optimization. For instance, the CVXTI algorithm exploits the divergence property of electric fields—manifested as a pronounced asymmetry in field intensity, with substantially greater strength on the side proximal to the electrode—to reformulate a non-convex problem into a convex optimization[14]. Furthermore, numerical computation-based methods[23–25] benefit from incorporating field distribution characteristics to reduce computational costs.

Here, we propose a quantitative computational approach from the perspective of the electric field to characterize its distribution properties. We have defined key concepts for tetrahedral meshes, including connected components, extremes, discrete ellipsoids, and the Local Moran’s index. Connected components are employed to delineate the spatial extent of electric fields exceeding a given threshold, while the discrete ellipsoid is used to quantify the spatial spread of the connected component. Data dimensionality is reduced through the identification of extremes and valid extrema, and the spatial clustering of these extremes is quantified using Local Moran’s index. Without presupposing stimulation targets, we analyze the spatial distribution of the TI electric field and produce comprehensive visualizations that delineate connected components and Local Moran’s index. Through this framework, we aim to determine effective stimulation targets in TI fields and quantify field focality.

## 2 Materials and methods

### 2.1 Head Model Construction

In this study, the head model used for the finite element simulation of the TI electric field is the example subject ‘ernie’ from simNIBS[26]. The head model was generated using the simNIBS charm pipeline (Fig. 1). Specifically, T1-weighted images, combined with T2-weighted images, were segmented into different types of tissues, including scalp, cranium, cerebrospinal fluid, gray matter, white matter, blood vessels, eyes, muscle and cavities. Subsequently, simNIBS converted the segmented images into a high-quality tetrahedral mesh and simultaneously generated the positions for the 10-10 electrode placement. Because the TI frequency satisfies the quasi-static approximation, allowing the displacement current to be neglected, the simulation used the tDCS boundary condition. The resulting electric field strength corresponds to the maximum amplitude of the interference envelope. All simulations were conducted using simNIBS, with the electrical conductivity parameters set to their default values.

**Fig. 1.**
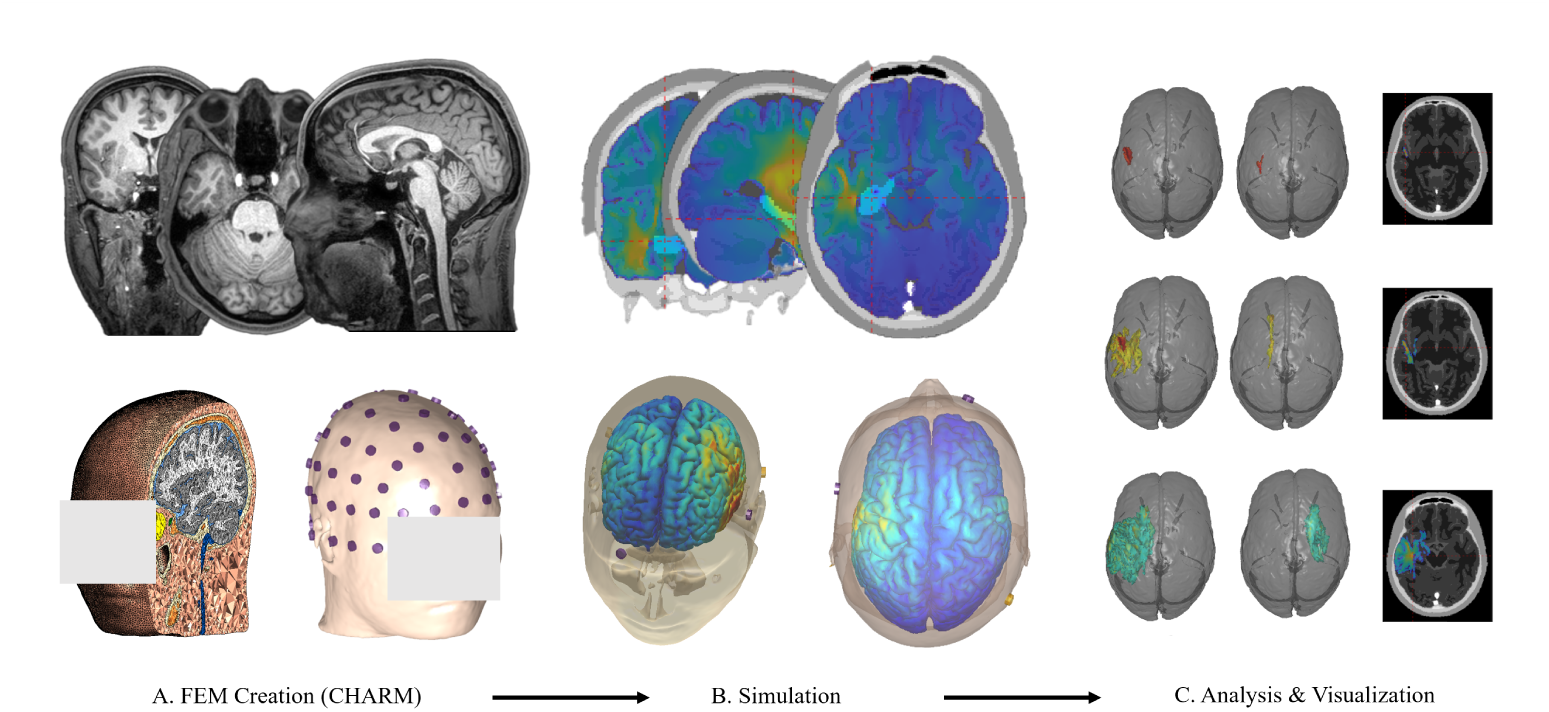
Workflow of the field-based analysis framework. (A) CHARM-based head modeling; (B) Electromagnetic field simulation and visualization; (C) Connected component analysis and visualization.

### 2.2 TI Envelope Electric Field Calculation

Different from tDCS and tACS, TI stimulation employs two sinusoidal signals with a small frequency offset. Since neurons are thought to be unresponsive to the highfrequency components but sensitive to the low-frequency envelope generated by signal superposition, the envelope amplitude, rather than the absolute peak value, is the primary determinant of the neural effect[4]. (1) can be used to compute the envelope electric field in the direction of the maximum envelope field intensity.

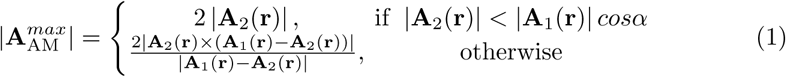

where **A**_1_(**r**) and **A**_2_(**r**) represent the vector amplitudes of the two interfering electric fields at location **r**, 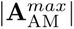 denotes the maximum envelope amplitude of the resultant signal, and *α* is the angle between the two field vectors **A**_1_(**r**) and **A**_2_(**r**), with *α < π/*2.

### 2.3 Neighborhood and Connected Component

In image processing, a connected component refers to an image region composed of pixels that share the same pixel value and are spatially adjacent to one another. Given the non-uniform spatial distribution of tetrahedral meshes, it is necessary to extend the concept of connected components. The extension of the connected component concept to non-uniform tetrahedral meshes begins with the definition of adjacency. Mesh elements are considered adjacent based on the sharing of a common node, edge, or face, thereby establishing a neighborhood. Building upon this definition, a connected component (CC) must satisfy two conditions (Fig. 2): (1) any two elements within it can be linked through a chain of neighboring elements, and (2) it is not a proper subset of any larger connected set of elements. In this work, adjacency is specifically defined as facial adjacency (i.e., sharing a common face); the precise mathematical definition is provided in Online Resource 1, Appendix A.

**Fig. 2.**
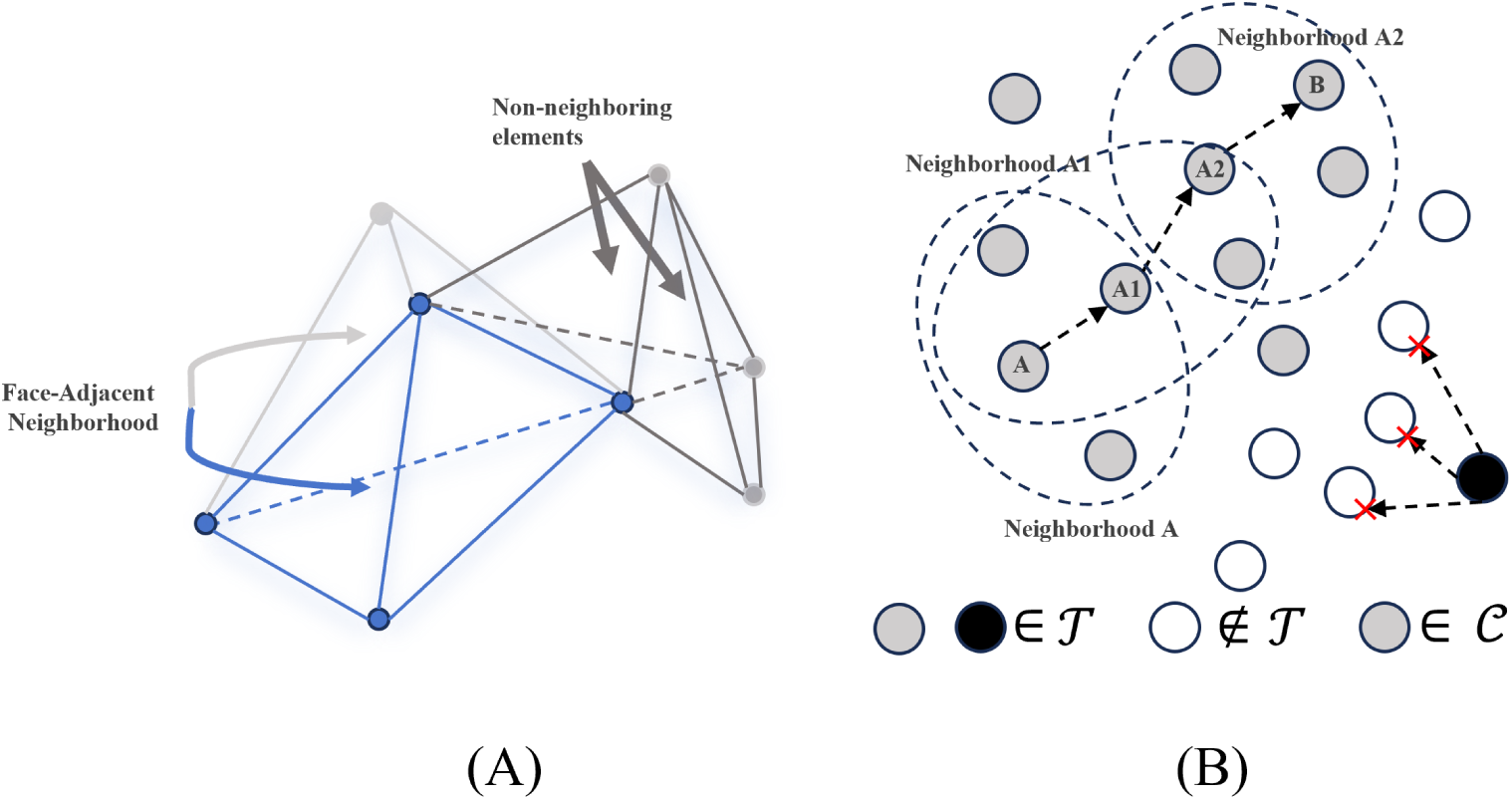
Schematic illustration of connected components. (A) Neighborhood: The light gray element shares a common face with the blue element, whereas the dark gray element shares only an edge or a node with the blue element. (B) Connected components: Both gray and black nodes belong to the set *T*. Each gray node can be linked via a chain of adjacent nodes, while the black nodes are separated by nodes not in *T*. Thus, the set of gray nodes forms one connected component, and the set of black nodes forms another.

### 2.4 Extrema and Valid Extremum

Given that the electric field permeates the entire brain across a computational mesh of millions of elements, it is neither practical nor necessary to analyze all data points. Specific analyses focus on distinct subsets: for example, focality is assessed using the distribution above the 50th percentile, while dose considerations target regions exceeding 0.2 V/m. Analogous to interpreting a topographic map (Fig. 3) by first identifying its highest peaks, we therefore introduce the concepts of extrema and valid extremum within the electric field to capture its most salient features for further analysis. An extremum is defined as a point where the electric field intensity is greater than that of all its neighboring points. A valid extremum is defined as an extremum that exceeds all other extrema within a defined spatial range, ensuring that only significant peaks are considered for subsequent analysis. The mathematical definitions of these concepts are provided in Online Resource 1, Appendix B.

**Fig. 3.**
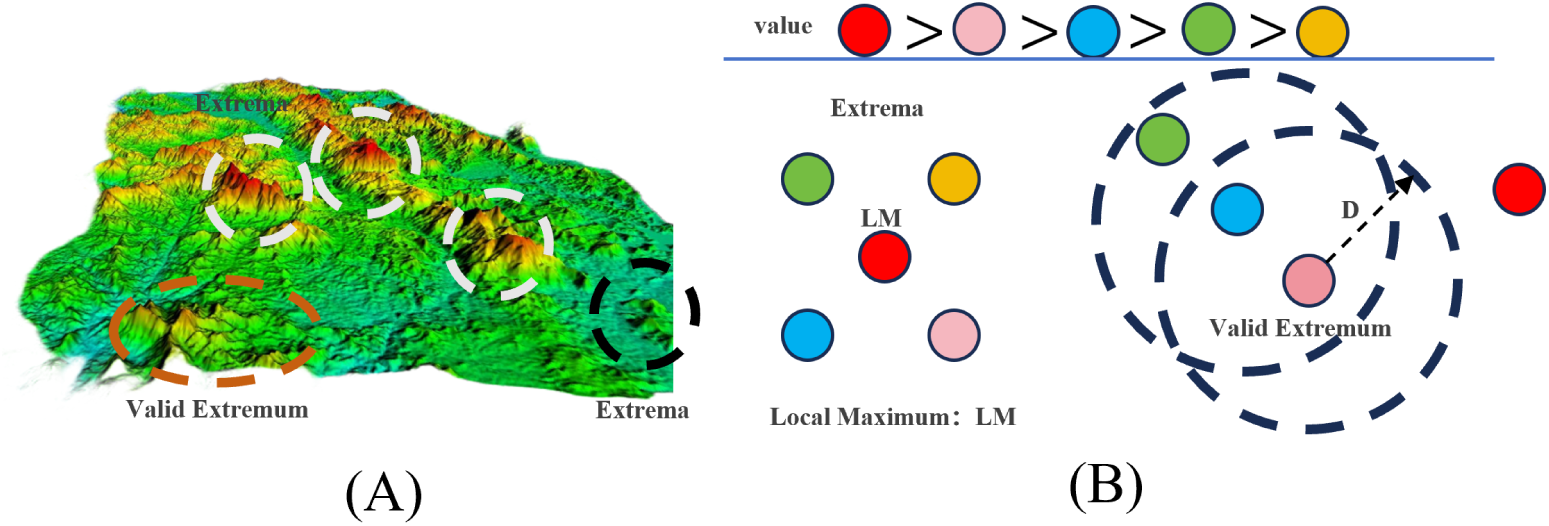
Schematic illustration of raw extremes and valid extremes.(A) Topographic map analogy: Peaks tend to draw attention regardless of their absolute intensity (black circles: low intensity; white circles: high intensity). When two peaks are in close proximity, the higher one becomes the focus of interest (red circle).(B) By analogy, an extremum is defined as the maximum value within its local neighborhood, while a valid extremum is defined as the maximum value within a prescribed distance D.

### 2.5 Field Metric

The most commonly used metric in existing literature is focality, typically defined as the cubic root of the total volume where the field strength exceeds 50% of its peak value. Other conventional metrics include the average electric field strength and field energy, the detailed definitions of which are given in [15]. A key limitation of these metrics lies in their global nature, primarily characterizing the overall field profile. They are often designed from a targeting-oriented perspective, serving as loss functions for optimization algorithms or as evaluative criteria for algorithmic performance. Consequently, such global metrics offer limited insight into the spatial distribution characteristics of the electric field.

As introduced earlier, connected components and extremes are defined to delineate focal regions and peak distributions, respectively. Building upon these, we aim to quantitatively describe the spatial location and extension of scalar fields, as well as to assess the degree and location of field focality. To this end, we introduce the concepts of the discrete ellipsoid and the Local Moran’s I index.

The discrete ellipsoid is conceptually inspired by the MATLAB function ***mesh get fieldpeaks and focality*** from the simNIBS toolkit, which calculates the center and dispersion of mesh element geometric centers exceeding a percentile threshold. In contrast to this point-set-based approach, our method defines dispersion directly on the mesh geometry: by analogy to the moment of inertia, we treat the electric field magnitude as a density distributed over the mesh, and the calculation is performed on individual connected components rather than on all elements meeting a threshold condition. The dispersion metric is formally defined as follows:

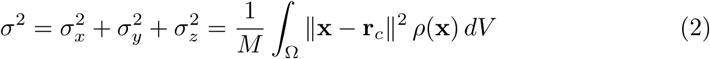

where Ω *⊂* ℝ^3^ denotes the region occupied by the connected component, *ρ*(**x**) is the electric field magnitude over Ω, 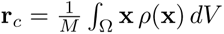 is the field-weighted centroid, and *M* = *∫*_Ω_*ρ*(**x**) *dV* is the total field mass. The quantities 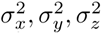 are the field-weighted variances along the *x, y, z* axes, respectively. A larger *σ*^2^ indicates a more widespread field, whereas a smaller value reflects tighter spatial focusing. The complete derivation is provided in Online Resource 1, Appendix C.

Relying solely on extremes is insufficient for characterizing the electric field distribution, primarily for three reasons: (1) the number of extremes remains excessively large; (2) extremes may include outliers introduced by computational artifacts or noise; and (3) the definition of extremes does not account for variations in mesh resolution or spatial proximity, leading to a higher density of extremes in regions with finer meshes. To address issue (1), we introduce valid extremes within a larger radius, which substantially reduces the number of candidate points. For issue (2), we incorporate Local Moran’s I to quantify the spatial clustering degree of extremes, mitigating the influence of outliers. To resolve issue (3), valid extremes are separately identified within a small radius and a large radius: the large-radius valid extremes serve as the central points for calculating Local Moran’s I, while the small-radius valid extremes provide the statistical sample for local aggregation assessment. This two-scale procedure explicitly accounts for local mesh density and spatial scale.

The Local Moran’s I index is formally defined as follows:

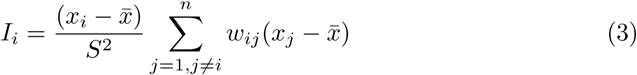

where *i* denotes the central spatial unit under evaluation (a large-radius valid extreme point), *j* indexes the other spatial units (small-radius valid extreme points), and *x_i_* is the electric field magnitude at location *i*. 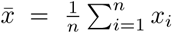 is the mean and 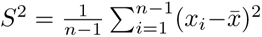 the sample variance of the attribute values, serving as normalization factors. The spatial weight *w_ij_* quantifies the strength of spatial connectivity between units *i* and *j* via a Gaussian kernel:

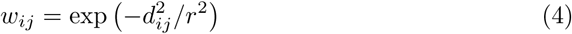

where *d_ij_* is the Euclidean distance between units *i* and *j*, and *r* controls the distance-based weighting.

### 2.6 Implementation Details and Parameters

The hippocampus represents a primary target in current TI stimulation studies, showing potential for enhancing memory and suppressing epileptic activity. As the montage proposed in previous literature [10] did not adhere to the standard 10-10 system, approximate 10-10 electrode positions (FT7-Fp2 and TP7-P8) were adopted in the simulations.

Because the TI field distribution scales linearly with the applied boundary conditions under the quasi-static approximation, understanding the relative spatial pattern of the field is more critical than its absolute magnitude. To extract connected components, we employed percentile-based thresholds, selecting six distinct levels: 95th, 99th, 99.9th percentile, and 50% of the 99.9th percentile, 75% of the 99.9th percentile, and an absolute value of 0.2 V/m. To prevent artificial fragmentation of connected components at the gray matter-CSF boundary, CSF was included in the computation of connected components. However, CSF contributions were excluded during subsequent metric calculation and visualization. Visualizations map field data onto the surface mesh, with each connected region colored by the electric field strength of the underlying tetrahedral elements. To improve clarity, the three principal axes of the discrete ellipsoid are scaled by a factor of three during rendering.

Valid extrema were computed separately for radii of 5 mm and 1 mm. The larger-radius extrema served as the central points for calculating the Local Moran’s I index, while the smaller-radius extrema provided the statistical sample for local aggregation assessment. A Gaussian kernel (r = 10 mm) was used as the spatial weight kernel in the Moran’s I computation.

During the preparation of this work, the authors used large language model-based tools to assist with language polishing and formatting. After using these tools, the authors reviewed and edited the content as needed and take full responsibility for the content of the published article.

## 3 Results

### 3.1 Connected Component

Based on the definition of connected components in Section 2.3, we calculated distributions under different thresholds. As shown in Fig. 4, the largest connected component (the main CC) accounted for most of the total connected-component volume across all thresholds, and the top few components consistently represented most of the volume. Table 1 shows that when the threshold was below 0.3 V/m, the main CC contributed over 70% of the total volume; as the threshold increased, this proportion decreased, indicating the presence of numerous scattered peaks. These scattered peaks are clearly visible in Fig. 5, where a large number of peaks are located near the brain surface— a pattern that may be related to CSF distribution, though computational artifacts could also contribute. It is noteworthy that the global maximum field value did not coincide with the main CC but appeared within smaller components. Based on the continuity assumption of the electric field, the maximum value within the main CC (0.702 V/m) may better represent the true peak intensity of the TI field.

**Fig. 4.**
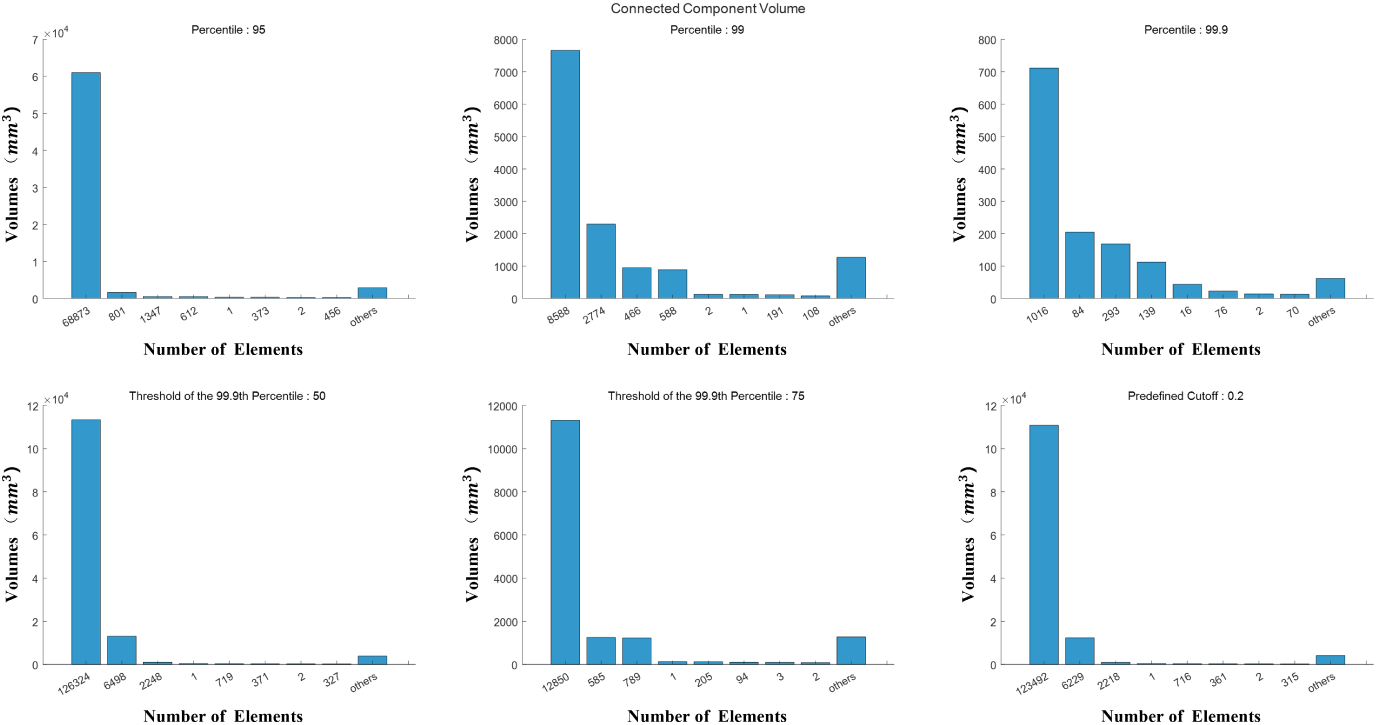
Volume of connected components of the TI field at different thresholds. For computational convenience, the volume here refers to the total volume of connected components with the same number of elements. All entries with a total volume greater than that of components containing one or two elements correspond to a single connected component.

**Fig. 5.**
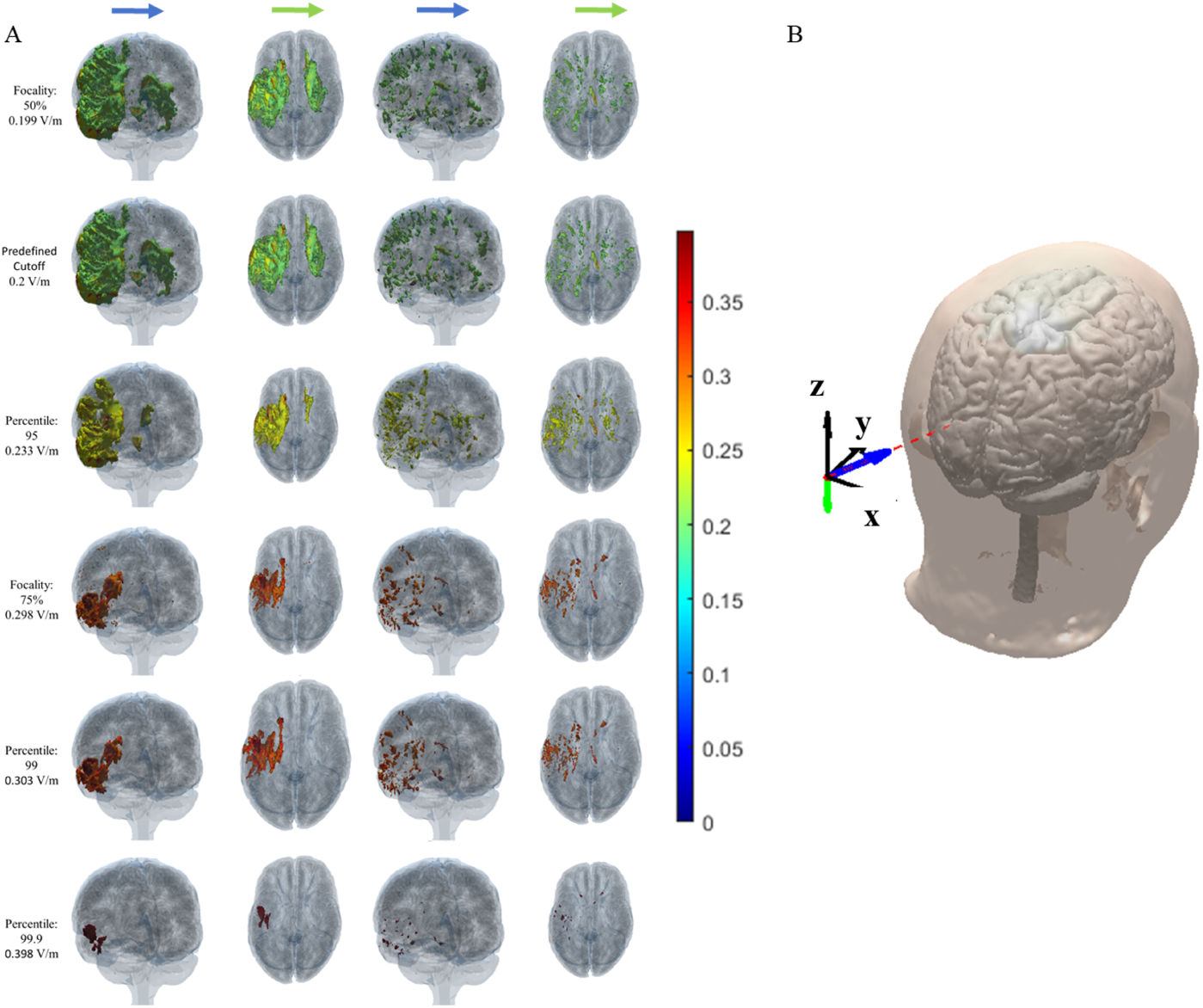
Connected components under different thresholds. (A) Connected components are arranged from top to bottom in ascending order of threshold intensity. The green and blue arrows indicate the viewing directions, which correspond to those shown in (B). From left to right, the first two columns display all connected components ranked among the top four by total volume (as defined in Fig. 4), while the last two columns show the remaining ones. The connected components are colored according to the electric field strength of the underlying tetrahedral elements, with warmer colors indicating higher field values. (B) Schematic diagram of viewing directions. The green arrow represents the topdown view, the blue arrow represents the lateral view, and the black arrows indicate the coordinate axes.

**Table 1.** Connected component analysis under different thresholds.

| Threshold Type | TI Field (V/m) | Main CC Volume (% Total) | Top 3 CC Vol (mm<sup>3</sup>) | Avg. TI (V/m) | Max. TI (V/m) | CC Vol. at Max (mm<sup>3</sup>) |
| --- | --- | --- | --- | --- | --- | --- |
| Percentile: 95 | 0.233 | 61032 (90.23%) | 1642<br>511<br>508 | 0.280 | 0.702 | 69 |
|  |  |  |  | 0.248 | 0.352 |  |
|  |  |  |  | 0.275 | 0.858 |  |
|  |  |  |  | 0.251 | 0.304 |  |
| Percentile: 99 | 0.303 | 7662 (56.70%) | 2293<br>951<br>888 | 0.359 | 0.702 | 39 |
|  |  |  |  | 0.329 | 0.457 |  |
|  |  |  |  | 0.316 | 0.349 |  |
|  |  |  |  | 0.322 | 0.426 |  |
| Percentile: 99.9 | 0.398 | 711 (52.63%) | 204<br>168<br>112 | 0.452 | 0.702 | 23 |
|  |  |  |  | 0.419 | 0.458 |  |
|  |  |  |  | 0.448 | 0.586 |  |
|  |  |  |  | 0.420 | 0.469 |  |
| Focality: 50% | 0.199 | 113391 (85.21%) | 13083<br>1025<br>358 | 0.251 | 0.702 | 85 |
|  |  |  |  | 0.219 | 0.352 |  |
|  |  |  |  | 0.244 | 0.858 |  |
|  |  |  |  | 0.252 | 0.698 |  |
| Focality: 75% | 0.298 | 11307 (72.38%) | 1252<br>1231<br>133 | 0.346 | 0.702 | 39 |
|  |  |  |  | 0.313 | 0.349 |  |
|  |  |  |  | 0.317 | 0.426 |  |
|  |  |  |  | 0.324 | 0.376 |  |
| Predefined Cutoff | 0.2 | 110845 (85.27%) | 12351<br>998<br>488 | 0.252 | 0.702 | 85 |
|  |  |  |  | 0.220 | 0.352 |  |
|  |  |  |  | 0.246 | 0.858 |  |
|  |  |  |  | 0.252 | 0.698 |  |

TI field values corresponding to each threshold: 95th (0.233 V/m), 99th (0.303 V/m), 99.9th (0.398 V/m), Focality: 50% (0.199 V/m), Focality: 75% (0.298 V/m), Predefined Cutoff (0.200 V/m).
CC: connected component; Vol: volume; Avg.: average; Max.: maximum; TI: the envelope electric field in the direction of the maximum envelope field intensity. Values for Avg. TI and Max. TI are presented as main CC / secondary CC / tertiary CC/quaternary CC.

Compared with conventional visualization methods (e.g., panels A and B in Fig. 1), the connected-component representation in Fig. 5 clearly reveals where the TI field is focused and its spatial extent above a given threshold. In contrast to the commonly used focality metric (typically defined at 50% of the peak value), Fig. 5 suggests that this metric may underestimate the spatial focality of TI stimulation: at the 75% focality threshold—only about one-third lower in field strength than the 50% level— the main CC volume is an order of magnitude smaller than at the 50% threshold (Table 1: 11,307 mm^3^ vs. 113,391 mm^3^). Therefore, the 75% focality threshold appears to better reflect the focused nature of the TI field.

The preceding analysis identifies three characteristic intensity levels for describing the TI field: the peak field, the 75% focality threshold, and the 0.2 V/m level. To systematically describe the spatial distribution of the field relative to these benchmarks, we introduce the following terms:

- **Peak distribution**: the set of connected components (CCs) where the electric field strength exceeds the 99.9th percentile.
- **Focal region**: the set of CCs where the electric field strength exceeds the 75% focality threshold.
- **Influential range**: the set of CCs where the electric field strength exceeds 0.2 V/m.

### 3.2 Discrete Ellipsoid

As shown in Fig. 5, the distribution of the TI electric field resembles a multilayered structure, where connected components corresponding to higher thresholds are enveloped by those of lower thresholds. This “layered” structure is geometrically irregular, exhibiting pronounced directional anisotropy. To characterize the directional extension of the TI field, we computed the spatial dispersion of connected components along three orthogonal axes (Section 2.5); the results are summarized in Table 2. As the threshold decreases, the growth of the connected component along the z axis is substantially greater than along the x axis. Given that the x axis points toward deeper brain regions (Fig. 5 (B)), this suggests that the TI field also exerts considerable influence on the cortical surface.

**Table 2.** Positional Discrete Ellipsoid(*ρ*(x) = 1)

|  | Percentile: 99.9%<br>Center (mm) | Focality: 75%<br>Center (mm) | Predefined Cutoff<br>Center (mm) |
| --- | --- | --- | --- |
| Volume | Axis ($\sigma_x^2, \sigma_y^2, \sigma_z^2$) | Axis ($\sigma_x^2, \sigma_y^2, \sigma_z^2$) | Axis ($\sigma_x^2, \sigma_y^2, \sigma_z^2$) |
| First | (-45.9, 20.9, 4.9) | (-40.0, 14.0, 1.8) | (-35.3, 13.7, 17.1) |
|  | (5.2, 26.2, 17.3) | (79.5, 117.0, 165.7) | (182.4, 258.8, 502.6) |
| Second | (-38.7, 14.2, -9.9) | (-19.2, 13.3, 20.0) | (26.2, 25.2, 20.6) |
|  | (1.0, 5.1, 10.4) | (5.8, 33.4, 14.4) | (32.9, 262.6, 105.8) |
| Third | (-37.5, 9.1, -1.4) | (-16.8, 34.1, 28.1) | (18.4, 55.8, -1.4) |
|  | (2.7, 24.5, 2.7) | (2.5, 305.2, 34.2) | (3.84, 50.6, 15.8) |
| Fourth | (-47.3, 5.8, -9.8) | (-31.3, 20.7, 12.9)* | (15.6, 16.8, 25.1)* |
|  | (1.3, 5.8, 2.8) | (335.4, 252.9, 497.4)* | (866.5, 476.5, 575.3)* |

Positional discrete ellipsoid parameters for the top four connected components ranked by total volume (as defined in Fig. 4).
\* indicates that there are multiple connected components with the same number of elements.

Notably, comparing Tables 2 and 3, the dispersion values obtained with and without field weighting show only minor differences, indicating that the spatial distribution of the electric field strength is largely consistent with the underlying mesh density. A plausible explanation is that the stimulation focus lies near the brain surface, where finer mesh elements are associated with lower field intensities and coarser elements in deeper regions with higher field values.

**Table 3.** Discrete Ellipsoid.

|  | Percentile: 99.9%<br>Center (mm) | Focality: 75%<br>Center (mm) | Predefined Cutoff<br>Center (mm) |
| --- | --- | --- | --- |
| Volume | Axis ($\sigma_x^2$) | Axis ($\sigma_x^2$) | Axis ($\sigma_x^2$) |
| First | (-45.9,20.9,5.0) | (-40.3,14.1,1.6) | (-35.4,13.9,16.1) |
|  | (5.1,25.6,17.0) | (75.9,113.1,158.3) | (179.3,250.8,487.0) |
| Second | (-38.7,14.2,-10.0) | (-19.3,13.3,20.0) | (26.2,25.6,20.7) |
|  | (1.0,5.0,10.2) | (5.8,33.1,14.2) | (32.6,263.4,105.2) |
| Third | (-37.5,9.1,-1.4) | (-16.8,34.0,28.1) | (18.3,55.7,-1.7) |
|  | (2.6,24.7,2.7) | (2.5,306.7,34.3) | (3.5,47.5,16.3) |
| Fourth | (-47.3,5.8,9.8) | (-31.3,20.7,12.8)* | (-15.6,16.8,25.1)* |
|  | (1.3,5.7,2.8) | (336.9,253.2,497.0)* | (866.2,477.8,576.1)* |

Discrete ellipsoid parameters for the top four connected components ranked by total volume (as defined in Fig. 4).
\* indicates that there are multiple connected components with the same number of elements.

As illustrated in Fig. 6, the discrete ellipsoid is constructed by taking the component’s centroid as the ellipsoid center and scaling its axes proportionally to the square root of the dispersion values (axes are magnified threefold for visualization). Compared to a scalar volume metric, this representation describes both the spatial location and anisotropic extension of the field distribution more clearly.

**Fig. 6.**
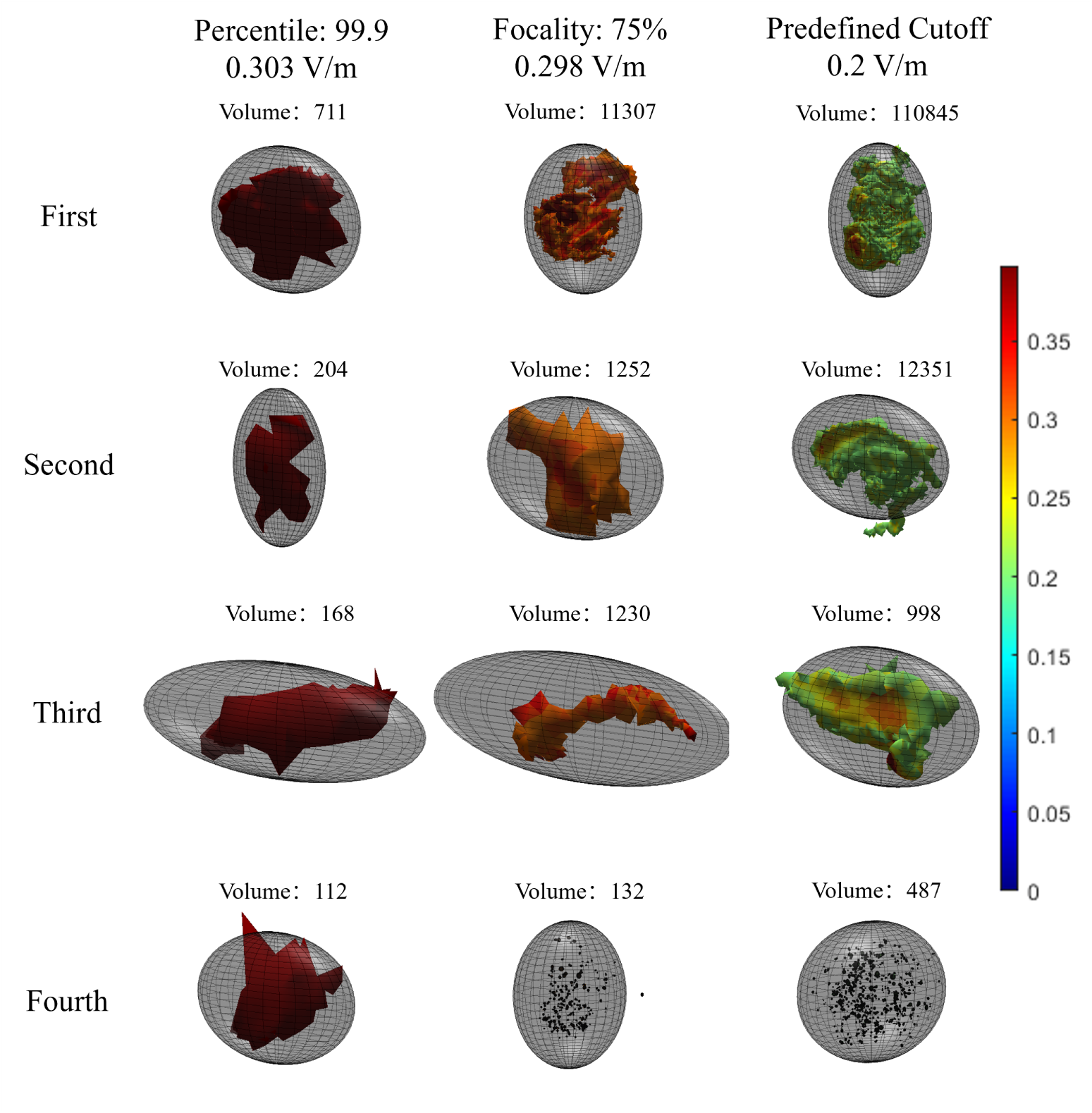
Visualization of positional discrete ellipsoids and connected components.

### 3.3 Extreme-based Dimensionality Reduction

Fig. 7 illustrates the data distribution and visualization of focality calculated using extreme values and the Local Moran’s I index. Panel A shows the distributions of the raw data, all extreme points, and valid extreme points within 5 mm and 1 mm radii. Panel B presents the Moran’s I values computed with the 5 mm valid extremes as centers and the 1 mm valid extremes as the local sample set. To enable intuitive visualization, the Moran’s I values were divided by 100 and shifted by adding the minimum value to ensure all results were positive (Panel C); this adjustment was applied solely for visualization and does not affect the statistical conclusions. All subsequent references to the Moran’s I index refer to this adjusted version. In Panel D, each 5 mm valid extreme is represented as a sphere, where color indicates the electric field magnitude and sphere diameter corresponds to the Local Moran’s I value; a larger sphere denotes a higher degree of local spatial clustering.

**Fig. 7.**
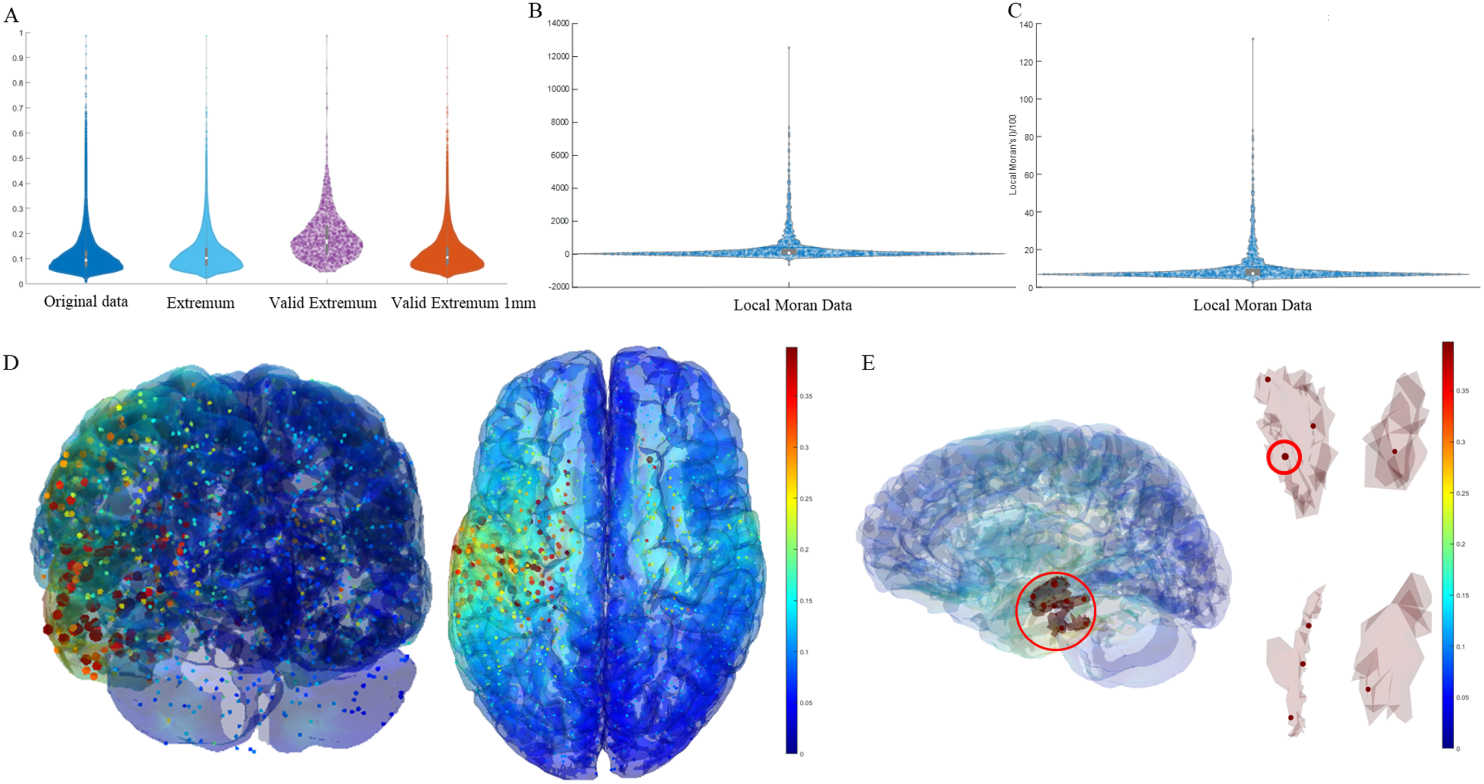
Distribution and visualization of extremes and Moran’s index. (A) Distribution of raw data, extremes, and valid extremes within radii of 5mm and 1mm. (B) Distribution of Moran’s index. (C) Distribution of the adjusted Moran’s index. (D) Visualization of Moran’s index. (E) Moran’s index within the peak distribution connected component.

Panel E analyzes the clustering distribution within connected components. The left side highlights the connected component corresponding to the peak distribution (marked by a red circle), while the right side enlarges four representative components for detailed inspection. Within the main connected component, the three strongest extremes are 0.70, 0.55, and 0.50 V/m, with Moran’s I indices of 132.07, 83.48, and 78.12, respectively. This is consistent with the peak connected component exhibiting the strongest spatial clustering. In contrast, the global maximum field value (0.99 V/m), which lies in a small peripheral component not listed in Table 1, yields a Moran’s I index of only 37.17, while the smallest Moran’s I within the peak CC is 68.72, corresponding to a field strength of 0.47 V/m. These results demonstrate that the Local Moran’s I index effectively mitigates outlier peaks and captures the intrinsic spatial aggregation of the electric field.

## 4 Discussion

This study proposes a topology- and mesh-aware computational framework for the characterization of transcranial interferential (TI) stimulation electric fields, overcoming key limitations of conventional global metrics through three interconnected analytical components: connected component analysis for delineating spatially continuous field regions, discrete ellipsoid modeling for quantifying directional spread and geometric centroids, and a two-scale extreme detection scheme combined with Local Moran’s I for identifying robust spatial clusters.

### 4.1 Where Is the Target? Beyond Uniqueness

Our findings indicate that the spatial extent of TI stimulation is considerably more complex than can be captured by a single target. Both the peak distribution and the focal region are distributed across multiple connected components located in distinct brain areas (Fig. 5). Nonetheless, the main connected component consistently accounts for more than 50% of the total volume, suggesting that it represents the primary region of neuromodulation. These observations imply that outcomes of clinical or animal studies should not be interpreted solely based on the intended target area; off-target contributions must also be considered.

Our results further demonstrate that the effective stimulation target in TI fields is more appropriately characterized as a spatially coherent volume rather than an isolated peak intensity point. Compared to the widely used 50% peak volume metric, the 75% peak volume threshold provides a more representative measure of spatial focality. In addition, connected component analysis clearly delineates the spatial extent of the electric field above a given threshold. Unlike approaches that rely exclusively on the global maximum, the Moran’s I index offers a more robust quantification of local clustering intensity. Accordingly, the location corresponding to the highest local Moran’s I value may be considered a candidate stimulation target—while recognizing that multiple such targets can coexist.

The proposed framework is inherently mesh-aware, incorporating the geometric and topological properties of the tetrahedral discretization to reduce sensitivity to mesh non-uniformity and enhance geometric consistency. A two-scale extreme detection strategy further improves robustness by integrating fine-scale local detail (1 mm radius) with broader regional context (5 mm radius), ensuring that the identified clustering patterns remain stable across spatial scales. This multi-scale, topology-driven approach naturally accommodates multiple targets, reinforcing the view that the “target” in TI stimulation is often not a singular locus but a set of spatially distinct regions.

### 4.2 What Does the Field Look Like? Multi-Metric Integration

Traditional focality metrics, such as the volume above 50% of the peak field, tend to overestimate the stimulated region and lack directional sensitivity. By integrating connected component volumetry with discrete ellipsoid orientation and Local Moran’s I clustering analysis, our framework provides a multi-faceted characterization of focality. The discrete ellipsoid captures anisotropic field spread, showing preferential extension, while Local Moran’s I identifies subregions of intense field clustering that are robust to numerical outliers. Moreover, the proposed quantitative descriptors can be readily employed to optimize stimulation parameters.

### 4.3 Visualization as an Interpretative Bridge

A distinctive contribution of this work is its emphasis on visually interpretable outputs. The framework generates layered component maps, discrete ellipsoid, and clustering maps of field extrema that directly translate complex field distributions into intuitive graphical representations. These visualizations aid in communicating spatial patterns and enable researchers to explore threshold-dependent field morphology and directional specificity—addressing a gap often left by purely quantitative, global metrics.

## 5 Conclusion

This study introduces a topology- and mesh-aware framework for analyzing transcranial interferential (TI) electric fields, integrating connected component analysis, discrete ellipsoid modeling, and Local Moran’s I. Applied to a publicly available template head model, the framework reveals that effective stimulation targets are better represented as spatially coherent volumes rather than isolated peak points. The discrete ellipsoid captures anisotropic field spread along the cortical surface, while Moran’s I identifies robust high-intensity clusters resilient to numerical outliers. Notably, multiple candidate targets may coexist across distinct brain regions, implying that experimental outcomes should not be attributed solely to intended target areas. The framework generates visually interpretable outputs—layered component maps, ellipsoids, and clustering landscapes—that bridge raw simulation data and physiological understanding. This work provides a coherent and visually intuitive toolkit for TI field analysis, advancing target localization, focality assessment, and parameter optimization in personalized neuromodulation.

## Supporting information

Online Resource 1

## Statements and Declarations

## Author Contributions

Tiandi Chen: Conceptualization, Methodology, Software, Formal analysis, Writing - original draft. Congcong Huo: Methodology, Investigation, Writing - review & editing. Guangjan Shao: Methodology, Investigation. Zhanhong Cao: Formal analysis, Writing - review & editing. Chao Li: Investigation, Writing - review & editing. Jizhong Liu: Supervision, Funding acquisition, Writing - review & editing. Zengyong Li: Conceptualization, Supervision, Funding acquisition, Writing - review & editing.

## Data Availability

The head model used in this study is the example subject ‘ernie’ distributed with the SimNIBS software package. The MATLAB analysis code is available from the corresponding author upon reasonable request.

## Ethics Approval

Not applicable. This is a computational simulation study using a publicly available template head model and involves no human or animal subjects.

## Consent to Participate

Not applicable.

## Consent for Publication

Not applicable.

