## Supplementary material for "Field-Based Characterization of Temporal Interference Stimulation: Beyond the Target-Centric Perspective": Online Resource 1

Mathematical definitions of neighborhood, connected component, extrema, and discrepancy

### 1 Appendix A: Neighborhood and Connected Component

In a tetrahedral mesh, two elements $T_i$ and $T_j$ are considered **neighbors** under a given adjacency condition. Common neighborhood types include:

- **Face-Adjacent Neighborhood (FAN)**:
  $T_i$ and $T_j$ share a common triangular face.
- **Edge-Adjacent Neighborhood (EAN)**:
  $T_i$ and $T_j$ share at least one common edge.
- **Vertex-Adjacent Neighborhood (VAN)**:
  $T_i$ and $T_j$ share at least one common vertex.

Here, $\mathcal{N}(T_i)$ denotes the **neighborhood** of $T_i$ under the specified adjacency rule.

Given a neighborhood relation $\mathcal{N}$ (e.g., FAN), a **connected domain** $\mathcal{C} \subseteq \mathcal{T}$ (where $\mathcal{T}$ is the set of all tetrahedra) is a **maximal connected subset** satisfying:

1. **Path Connectivity** :
$$\forall T_i, T_j \in \mathcal{C}, \exists \{T_{i_1}, T_{i_2}, \ldots, T_{i_k}\} \subseteq \mathcal{C}, \text{such that}$$
$$T_{i_1} = T_i, T_{i_k} = T_j, \text{ and } T_{i_m} \in \mathcal{N}(T_{i_{m+1}})$$
$$\text{for } m = 1, \ldots, k-1.$$
   (Any two tetrahedra in $\mathcal{C}$ are connected via a chain of neighboring elements.)
2. **Maximality**:
$$\nexists T \in \mathcal{T} \setminus \mathcal{C} \text{ such that } \mathcal{C} \cup \{T\} \text{ remains connected under } \mathcal{N}.$$
   (No additional tetrahedron can be included without breaking connectivity.)

### 2 Appendix B: Extremum

#### 2.1 Extrema

Let $\mathcal{T} = \{T_i\}$ be a tetrahedral mesh where each element $T_i$ carries a scalar value $f(T_i)$. Given a neighborhood relation $\mathcal{N}(T_i)$ (e.g., FAN/EAN/VAN):

##### Local Maximum

A tetrahedron $T^*$ is a **strict local maximum** if:

$$f(T^*) > f(T_j) \quad \forall T_j \in \mathcal{N}(T^*)$$

**Non-strict** version: $f(T^*) \geq f(T_j)$.

##### Local Minimum

A tetrahedron $T_*$ is a **strict local minimum** if:

$$f(T_*) < f(T_j) \quad \forall T_j \in \mathcal{N}(T_*)$$

**Non-strict** version: $f(T_*) \leq f(T_j)$.

#### 2.2 Definition: Valid Extremum Point

For detected extremum points (maxima or minima), we define their **validity** as follows:

##### Valid Local Maximum

A strict local maximum element $T^*$ (satisfying $f(T^*) > f(T_j), \forall T_j \in \mathcal{N}(T^*)$) is called a **valid local maximum** if:
Within a spherical region $B_R(\mathbf{x}^*)$ of radius $R$ centered at $T^*$'s geometric centroid $\mathbf{x}^*$, **no other strict local maximum** $T'$ **exists such that** $f(T') \geq f(T^*)$.
Formally:

$$\nexists T' \in \mathcal{T} \cap B_R(\mathbf{x}^*), \quad \text{s.t.} \quad \begin{cases} f(T') > f(T_j), \forall T_j \in \mathcal{N}(T'), \\ f(T') \geq f(T^*). \end{cases}$$

##### Valid Local Minimum

Similarly, a strict local minimum element $T_*$ is **valid** if no other strict local minimum $T'$ within radius $R$ satisfies $f(T') \leq f(T_*)$:

$$\nexists T' \in \mathcal{T} \cap B_R(\mathbf{x}_*), \quad \text{s.t.} \quad \begin{cases} f(T') < f(T_j), \forall T_j \in \mathcal{N}(T'), \\ f(T') \leq f(T_*). \end{cases}$$

### 3 Appendix C: Discrepancy

The discrepancy ($\sigma^2$) is defined as:

$$\sigma^2 = \frac{1}{M} \int_\Omega \|\mathbf{x} - \mathbf{r}_c\|^2 \rho(\mathbf{x})\, dV \tag{1}$$

where $\mathbf{x}$ is any point in the object, $\mathbf{r}_c$ is the geometric center, $\rho(\mathbf{x})$ is the field value, and $M = \int_\Omega \rho(\mathbf{x})\, dV$ is the total field value.

#### 3.1 Discretization

The mesh consists of $N$ tetrahedral elements. Each element $e$ has volume $V_e$, constant field value $\rho_e$, and total field $m_e = \rho_e V_e$. Its geometric centroid is $\mathbf{r}_e$. The global field center is $\mathbf{r}_c = \frac{1}{M} \sum_e m_e \mathbf{r}_e$.

#### 3.2 Discrete Formula

Using a local coordinate $\boldsymbol{\xi} = \mathbf{x} - \mathbf{r}_e$ within each element, the squared distance expands as:

$$\|\mathbf{x} - \mathbf{r}_c\|^2 = \|\mathbf{r}_e - \mathbf{r}_c\|^2 + 2(\mathbf{r}_e - \mathbf{r}_c) \cdot \boldsymbol{\xi} + \|\boldsymbol{\xi}\|^2.$$

Since $\int_{\Omega_e} \boldsymbol{\xi}\, dV = \mathbf{0}$, the linear term vanishes. Summing over elements gives:

$$\sigma^2 = \frac{1}{M} \sum_{e=1}^{N} m_e \left( \|\mathbf{r}_e - \mathbf{r}_c\|^2 + \frac{1}{V_e} \int_{\Omega_e} \|\boldsymbol{\xi}\|^2\, dV \right).$$

#### 3.3 Inertia Moment of a Tetrahedron

For a tetrahedron with vertices $\mathbf{X}_0, \mathbf{X}_1, \mathbf{X}_2, \mathbf{X}_3$ and centroid $\mathbf{r}_e = \frac{1}{4} \sum_{i=0}^{3} \mathbf{X}_i$, the integral of $\|\boldsymbol{\xi}\|^2$ over its volume can be derived using barycentric coordinates and standard simplex integrals. The result is:

$$\int_{\Omega_e} \|\boldsymbol{\xi}\|^2\, dV = \frac{V_e}{20} \sum_{i=0}^{3} \|\mathbf{X}_i - \mathbf{r}_e\|^2.$$

#### 3.4 Final Expression

Substituting this closed form, the discrete discrepancy becomes:

$$\boxed{\sigma^2 = \frac{1}{M} \sum_{e=1}^{N} m_e \left( \|\mathbf{r}_e - \mathbf{r}_c\|^2 + \frac{1}{20} \sum_{i=0}^{3} \|\mathbf{X}_i^{(e)} - \mathbf{r}_e\|^2 \right).}$$
